# Predicting local tissue mechanics using immunohistochemistry

**DOI:** 10.1101/358119

**Authors:** David E. Koser, Emad Moeendarbary, Stefanie Kuerten, Kristian Franze

## Abstract

Local tissue stiffness provides an important signal to which cells respond *in vivo.* However, assessing tissue mechanics is currently challenging and requires sophisticated technology. We here developed a model quantitatively predicting nervous tissue stiffness heterogeneities at cellular resolution based on cell density, myelin and GFAP fluorescence intensities. These histological parameters were identified by a correlation analysis of atomic force microscopy-based elasticity maps of spinal cord sections and immunohistochemical stainings. Our model provides a simple tool to estimate local stiffness distributions in nervous tissue, and it can easily be expanded to other tissue types, thus paving the way for studies of the role of mechanical signals in development and pathology.

## Introduction

Cells sense and respond to the stiffness of their environment. In the central nervous system (CNS), for example, neuronal growth is regulated by local tissue stiffness in the developing brain of *Xenopus in vivo* (1). Astrocytes and microglial cells, on the other hand, which are glial cells involved in a multitude of physiological and pathological processes in the CNS, increase the expression in inflammatory markers when exposed to inappropriate mechanical signals *in vivo* (2).

Healthy CNS tissue is mechanically highly heterogeneous (1, 3-8), and its mechanical properties change during development (1, 5) and a variety of pathological processes (8, 9). Such spatial or temporal alterations in tissue stiffness will impact the behaviour of cells during development and disease (10, 11). Thus, knowledge of local tissue stiffness distributions is crucial to understand mechanobiological processes.

While a number of techniques have recently been developed to measure tissue mechanics at cellular or sub-cellular resolution, including atomic force microscopy (AFM) (12), Brillouin microscopy (13), and magnetically responsive ferrofluid microdroplets (14), measurements are technically challenging, and access to those techniques is often limited. To overcome these obstacles, we here developed an empirical model predicting local tissue stiffness at a cellular length scale solely based on immunofluorescence imaging.

## Materials and Methods

### Animals

All animal experiments were performed in accordance with regulations issued by the Home Office of the United Kingdom under the Animals (Scientific Procedures) Act of 1986. Five female SJL mice (4-7 weeks old) were obtained from Janvier Labs (Saint Berthevin Cedex, France) and housed in open topped cages in a conventional animal facility. Mice were kefBatoulpt in groups of 2-3 animals per cage with woodchip bedding. Pelleted food and water was provided *ad libitum* and mice were kept in an artificial 12/12 hours dark/light cycle. As these mice served as controls in a study on experimental autoimmune encephalomyelitis (Koser et al., in preparation), they were injected with a mixture of PBS and Complete Freund’s adjuvant (complete injection volume: 200 μL) subcutaneously in the flanks 15-62 days before the measurements (15).

### Acute slice preparation

#### Solutions

All chemicals were obtained from Sigma-Aldrich (Sigma-Aldrich Company Ltd., Gillingham, UK) unless stated differently. The slicing and measuring artificial cerebrospinal fluids (s-aCSF and m-aCSF, respectively) were composed as described in Mitra and Brownstone (16). Both solutions were prepared freshly before each experiment and constantly bubbled with 95% O2/ 5% CO2. The resulting pH was ~7.3.

#### Dissection

Mice were sacrificed by a preserved method of cervical dislocation. Dissection was done as previously published (16). Briefly, mice were fixed on an ice-cold aluminium-wrapped polystyrene pad and then eviscerated in order to expose the vertebral column ventrally. The abdomen was washed and filled with ice-cold s-aCSF, which was exchanged every two minutes. The time between cervical dislocation and first contact with s-aCSF was kept under ~8 minutes. Next, the spinal cord was exposed and the ventral and dorsal roots and part of the meninges were cut craniocaudally, while the cord was gently lifted up at the cranial end of the exposed area. The spinal cord was transferred to a s-aCSF-filled petri dish, and the dura mater and the remaining roots were carefully removed under a stereomicroscope while s-aCSF was exchanged every two minutes. An aluminium-foil cube (2×2×2 cm^3^) was half filled with 4% low melting point agarose in 0.1 M PBS solution at 37 °C. The lumbar enlargement of the spinal cord was placed inside the cube, and the cube was carefully filled up with low melting point agarose. The cube was directly placed on ice to solidify the agarose. This procedure was carried out within ~30 minutes after sacrificing.

#### Slice preparation

Slice preparation was done as previously described (6). Briefly, the spinal cord-containing agarose block was glued onto a vibratome (VT1000 S, Leica Microsystems Ltd, Milton Keynes, UK) plate, and transverse spinal cord slices were obtained. The plate was submerged in ice-cold s-aCSF bubbled with 95% O2/ 5% CO2. The vibratory amplitude was set to 1 mm with a frequency of 100 Hz and the forward speed of ~40 μm/s. Eight slices with the thickness of 500 μm were obtained from the lumbar region of the spinal cord. Four odd-numbered slices were directly fixed in 4% PFA for ~4 h and later used for immunohistochemistry. The remaining four even-numbered slices were transferred to a custom-built storage container, which was filled with m-aCSF and bubbled with 95% O2/ 5% CO2. One or two of these slices were used for AFM measurements and the corresponding fixed slices were used for immunohistochemistry. The second slice was only measured if the measurement could be finished within less than ~8 h after euthanasia. Slices were transferred to petri dishes coated with Cell-Tak (BD Cell-Tak Cell and Tissue Adhesive, BD Biosciences, Oxford, UK) and covered with m-aCSF, which was constantly exchanged by bubbled m-aCSF through a custom-built flow system with a flow rate of ~0.33 mL/min. Before the AFM measurement started, the slice was left to recover in m-aCSF for at least 15 min.

### Atomic force microscopy (AFM)

All AFM measurements were done within ~8 h *ex vivo*, during which the CNS tissue stays viable and its mechanical properties do not change (4, 6, 17). Measurements were performed using a JPK Nanowizard Cellhesion 200 AFM (JPK Instruments AG, Berlin, Germany), which was set up on an inverted optical microscope (Axio Observer.A1, Carl Zeiss Ltd., UK). The spring constant of the tipless silicon cantilevers (Arrow-TL1, NanoWorld, Neuchatel, Switzerland) was determined via the thermal noise method (18) and cantilevers with a spring constant between 0.01 and 0.02 N/m were selected. Monodisperse polystyrene beads (d =37.28 ± 0.34 μm; microParticles GmbH, Berlin, Germany) were glued to the tip of the cantilevers and used as probes. Slices in Cell-Tak-coated petri dishes were transferred to the AFM-setup, and images of the sections were taken with a CCD camera (The Imaging Source, Bremen, Germany). The ventral half of each spinal cord slice was selected for AFM measurement. Force-distance curves were automatically taken every 50 μm in a raster scan using a custom-written script with a maximum force of 7 nN and an approach speed of 10 μm/s. Images of the upper right and lower left corners of the selected rectangular region were taken to match the measured region with the image of the slice (6).

Force-distance curves were analysed as described previously (6). Briefly, the curves were analysed for the full indentation depth at F = 7 nN and fitted with the Hertz model to extract the apparent elastic modulus *K*, which is a measure of tissue stiffness. The stiffness values were mapped onto the image of the slice, resulting in elasticity maps as shown in Fig. 1A. To extract region-specific (white and grey matter) stiffness values, ROIs were segmented by a custom-written MATLAB routine.

**Figure 1:**
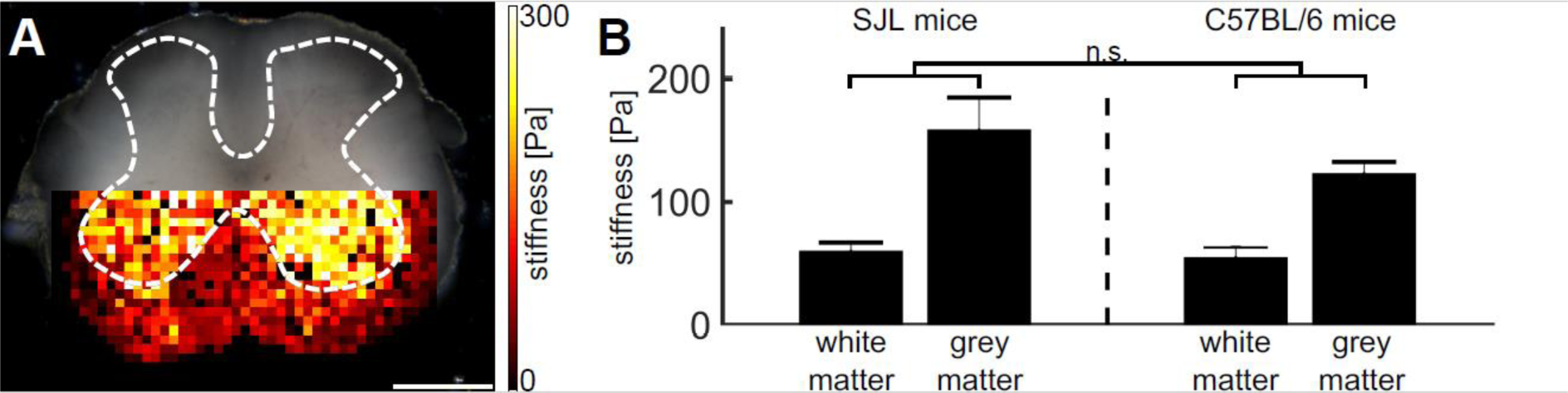
Mechanical heterogeneities within the mouse spinal cord. (**A**) Representative elasticity map of the ventral half of a transverse spinal cord section obtained by AFM, displaying clear local heterogeneities that are correlated with white and grey matter areas. The elasticity map is overlaid on a bright-field image of the spinal cord section. The border between white and grey matter is indicated by a white dashed line. **(B)** White matter was significantly softer than grey matter in the tested female SJL mice, and the results were comparable to our previous studies in C57Bl/6 mice (6). Scale bar: 500 μm.

### Fluorescence imaging

Fixed spinal cord slices were paraffin-embedded using an automatic tissue processor (Leica TP 1020, Leica Biosystems Nussloch GmbH, Nussloch, Germany) or a standard manual paraffin embedding protocol. Slices with ~5 μm thickness were cut using either a Leica SM 2010R or a Leica SM 200R microtome (Leica Biosystems Nussloch GmbH, Nussloch, Germany) and mounted on microscope slides (Superfrost Plus, Thermo Fisher Scientific Inc., Waltham, MA, USA). Slices were deparaffinized by incubation in xylol twice for 10 min, and subsequently in decreasing percentages of ethanol (100%, 100%, 96%, 80%, and 70%) for 5 min, and finally washed in distilled water for 5 min.

Antigen retrieval was performed using 0.1 M citrate buffer (pH 6.0) by boiling the sections in citrate buffer. Sections were then washed with 0.1 M PBS, and blocked for 1 h with blocking reagent (MOM kit, Vector Laboratories, CA, USA) in case of MBP staining or otherwise for 2 h in 5% normal goat serum (NGS). Primary antibodies were incubated for 30 min (MBP; 1:500, cat. no. SMI-99P, Covance, NJ, USA) in diluent solution (MOM kit) or at 4 °C overnight (GFAP; 1:3000, cat. no. G3893, Sigma-Aldrich, Germany, and Iba1; 1:200, cat. no. 019-19741, Wako, Germany) in 1% NGS, 0.2% bovine serum albumin (BSA). Sections were washed in PBS and incubated with the secondary antibody for 30 minutes (MBP; biotin-conjugated goat anti-mouse IgG, 1:250, Vector Laboratories, CA, USA) or for 1 h (GFAP and Iba1; goat-anti-mouse Cy5 and goat-anti-rabbit Cy5, respectively, 1:400 for both, Dianova, Germany). Unbound antibodies were washed off using PBS, and sections incubated for 30 min in NeutrAvidin Dylight 550 (1:400, Pierce, IL, USA) in case of MBP, washed again, and in any case incubated for 10 min with 4’,6-diamidino-2-phenylindole (DAPI; 1:5000). Finally, sections were washed and mounted.

Imaging was performed using a laser scanning confocal microscope (Nikon A1, Nikon, Germany) using a 20x objective (Plan Apo, NA=0.8). Images were taken in a raster scan with 15 - 20% overlap and subsequently stitched. At each position, a z-stack consisting of 3-7 images was acquired with a step size of 1-2 μm. ImageJ (National Institutes of Health, Bethesda, MD, USA) was used to process the stacked images; maximum intensity projections were used for further analysis. First, cell density was evaluated by semi-automatic thresholding of the DAPI images, similar to previous reports (19, 20). Briefly, images were processed by discarding intensity values lower than the mean + 1.96 times the standard deviation of the background intensity distribution (corresponding to a 97.5% confidence interval). A manual analysis of more than 100 DAPI-labelled cells revealed minimum nuclear areas of ~4.8 μm^2^ (corresponding to 50 pixels); thus connected areas with less than 4.8 μm^2^ were discarded from further analysis. Next, the area was binarised, 8-connectivity was eliminated by diagonal filling and a majority criterion was used to fill pixel gaps. Furthermore, connected ‘holes’ of 9 pixels within each area were filled in any case and connected ‘holes’ of 10-16 pixels within the area were filled if the eccentricity of the area was below 0.6. A binarised image was created as an output. By using the described method a good segmentation of the area taken up by cell nuclei was achievable. Note that in some cases the area taken up by cell nuclei in grey matter was underestimated as nuclear areas of large motor neurons may not be correctly detected. Next, MBP, Iba1, and GFAP intensity images as well as binarised cell density images were registered with the corresponding wide-field image taken during the AFM measurement using affine transformation and cross-correlation. The registered images of cell density and normalized fluorescence intensity were downscaled to match the 50 μm resolution of the stiffness maps, resulting in the normalised matrices *P*_*nuc*_*iei, ¡mbp, Igfap*, and //&_α1_. This approach allowed accurate investigation of the correlation between the measured histological parameters and local tissue stiffness within one sample. For better visualization of the immunohistochemical images shown in Fig. 2, the brightness/contrast was adjusted, and the ventral roots of the spinal cords were covered.

**Figure 2:**
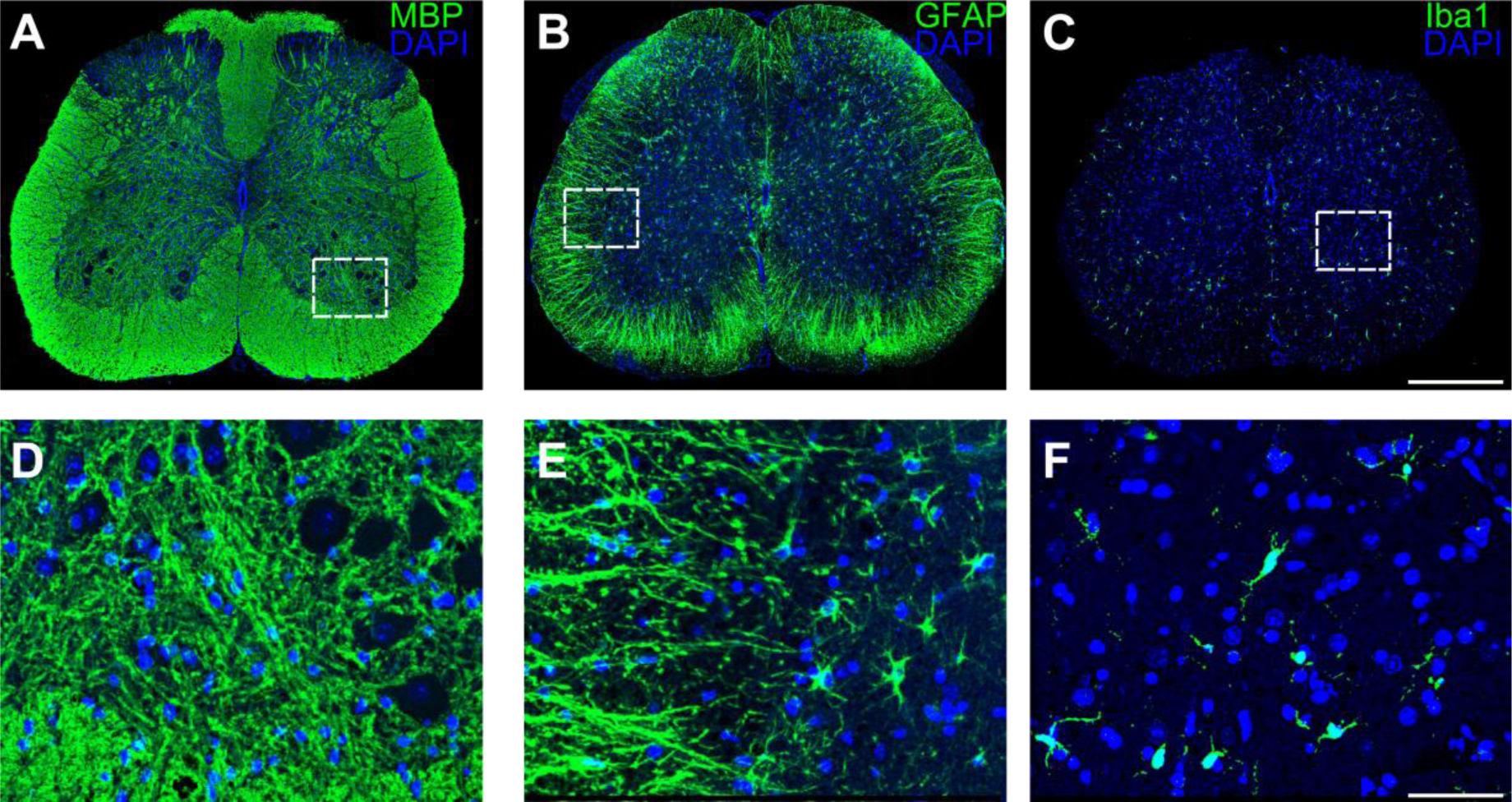
Structural heterogeneities within the mouse spinal cord. (**A-C**)Immunohistochemistry of the mouse spinal cord. Representative staining for **(A)** myelin (MBP), **(B)** astrocytes (GFAP), and **(C)** microglia/macrophages (Iba1), all combined with staining of cell nuclei (DAPI). **(D-F)** Higher magnifications of the regions within the white dashed rectangles shown in **(A-C)**. Scale bar for **(A-C)**: 500 μm. Scale bar for **(D-F)**: 75 μm.

### Multiple linear regression analysis

For the generation and evaluation of linear models, the *stepwiselm, stepwisefit* and *regress* MATLAB functions were used. Two approaches were employed: In the first, we started with a constant model and added any parameters that statistically increased the predictive power of the model (evaluated based on the change in the sum of squared errors; p-values from F-tests); this approach was termed minimal linear model. In the second approach, we started with including all parameters and then removed parameters and/or added interacting terms according to their significance; this approach was termed full linear model.

### Quantification and statistical analysis

All data and statistical analyses as well as plotting were performed in MATLAB (Mathworks). Data were tested for normal distribution by the Lilliefors test. In case two groups were compared, significance was tested either with Student’s t-test (in case of normal distribution) or Mann-Whitney *U* test (in case of non-normal distribution, or ordinal data). If more than two groups were compared, ANOVA or Kruskal-Wallis tests were used, in combination with the Bonferroni-corrected *post hoc* tests. Two-tailed tests were used throughout the statistical analysis. In the text we mention either the mean ± SEM or the median. Pearson’s correlation coefficients (ρ) are reported for linear models.

## Results

Stiffness maps of the ventral half of acute transverse spinal cord slices from the lumbar enlargement of female SJL mice were obtained using AFM with an in-plane resolution of 50 μm (Fig. 1A, approximately 900 measurements on each spinal cord tissue). In all slices *(N* = 5 slices from *5* animals), grey matter was significantly stiffer than white matter (p < 10^−6^ in all slices, Mann-Whitney *U* tests). The average median apparent elastic moduli *K* for grey and white matter were *K* = 159 ± 26 Pa and 60 ± 7 Pa, respectively, which were statistically similar to values measured in our previous study (K = 123 ± 9 Pa and 55 ± 9 Pa, respectively) in a different mouse strain (6), confirming the robustness and consistency of AFM experiments (Fig. 1B; p > 0.5, Bonferroni-corrected *post hoc* tests).

We obtained spinal cord slices of the same animals (see *Material and Methods)* for histological staining. As previous studies showed that cell densities (1, 6, 21), myelination (22-24), and glial activity (8, 25) influence tissue mechanics, we stained for DAPI (cell nuclei), MBP (myelin), GFAP (astrocytes), and Iba1 (microglia/macrophages) (Fig. 2). The cell density (approximated by DAPI quantification; SI Fig. 1) and the normalized fluorescence intensities of MPB, GFAP, and Iba1 were used as a readout. We then applied a multiple linear regression analysis (26) to identify a system of histological parameters (see *Material and Methods*) reproducing the identified mechanical heterogeneities on a 50μm scale (Fig. 1A). Both a minimal and a full approach (see *Material and Methods)* resulted in the same linear model containing the cell density (p < 10^−5^, F-test), MBP intensity (p < 10^−26^, F-test), GFAP intensity (p < 10^−13^, F-test), and an interaction term between MBP and GFAP (p < 10^−8^, F-test):

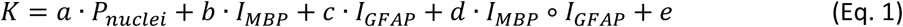

with *K* being an apparent elastic modulus (i.e., a measure of mechanical stiffness), *P_nuc_i_e_i* the estimated density of cell nuclei, *I* the fluorescence intensities, *a-e* tissue-specific constants (see Table 1), and ° the Hadamard product.

**Table 1:**
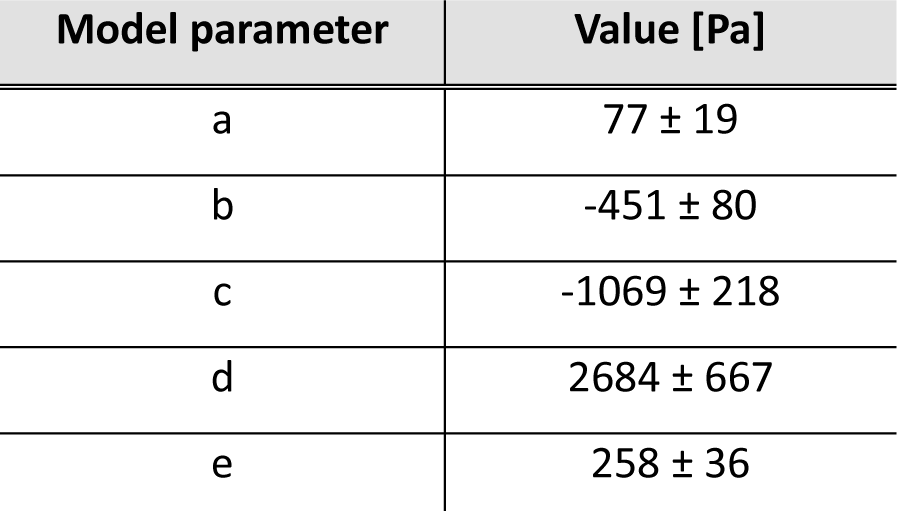
Model parameters predicting spinal cord tissue stiffness at a 50 μη scale.

In agreement with our previous studies (1, 6, 21), we observed a positive correlation between tissue stiffness and cell density. MBP and GFAP intensities, on the other hand, correlated negatively with tissue stiffness, and their interaction term positively. Microglia/macrophages (stained by Iba1), which are quiescent in healthy neural tissue, did not influence the observed mechanical heterogeneities (p > 0.2, F-test).

The above described model was not only sufficient to recapitulate and even predict the overall mechanical heterogeneities within a single spinal cord section (Fig. 3B; ρ = 0.66), but for all tested spinal cord sections (Fig. 4A,B; ρ = 0.52 ± 0.04). The model performed very accurately, with a model error - defined as the relative over-/underestimation of the measured stiffness - of 8 ± 2% (Fig. 4C). Furthermore, adding a Gaussian noise term, to account for measurement uncertainties, yielded stiffness maps closely resembling the maps experimentally obtained by AFM (Fig. 3C), indicating that the model accurately predicts local tissue stiffness of transverse spinal cord sections.

**Figure 3:**
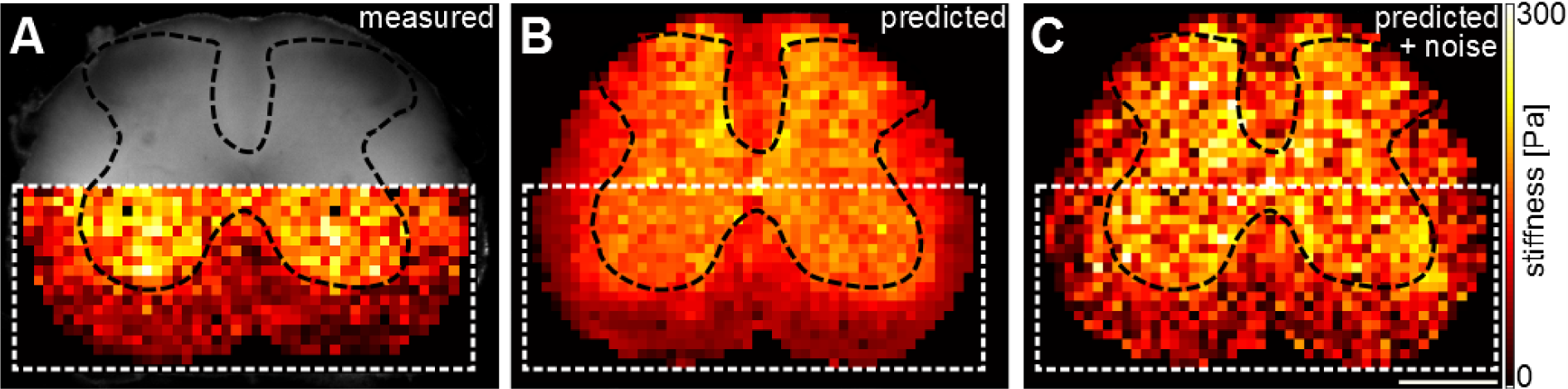
Mechanical heterogeneities are predicted by the underlying cellular structures. (**A)** Representative elasticity map of the ventral half of a transverse spinal cord section assessed by AFM indentation measurements. The border between white and grey matter is indicated by a black dashed line. **(B)** Corresponding elasticity map reconstructed from the linear model (Eq. 1, Tab. 1). **(C)** Adding a Gaussian noise term to the model results in a map closely resembling the original stiffness map shown in (A). White dashed boxes in (B, C) highlight the area measured by AFM shown in (A). Scale bar: 500 μm.

**Figure 4:**
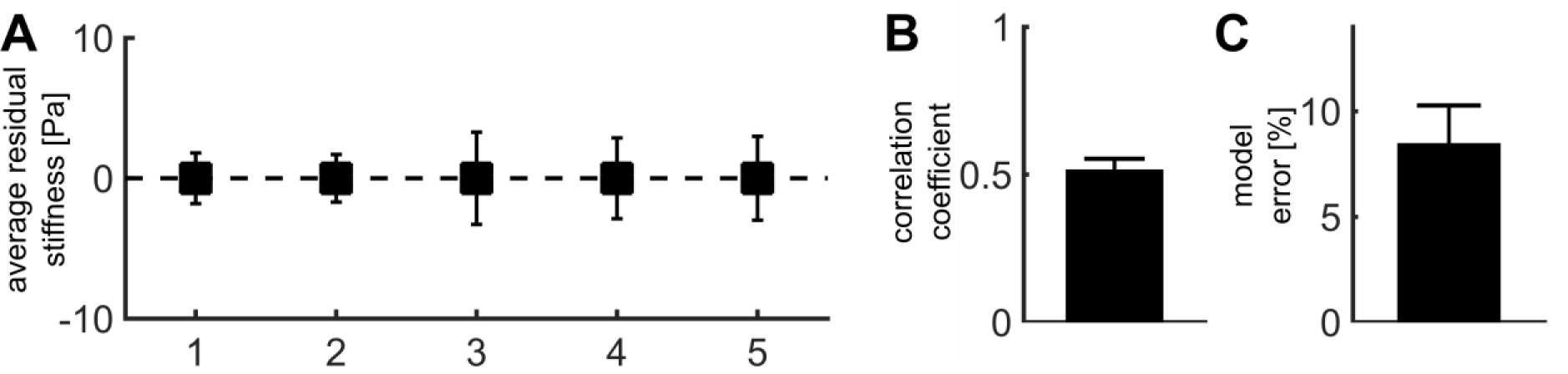
Accuracy of the model predicting local spinal cord stiffness. (**A**) Average stiffness difference between model and measurements (defined as residual stiffness) is shown for each of the 5 tested spinal cord slices. **(B)** Correlation coefficient between model sand measurements. **(C)** Relative over/underestimation of measured stiffness by the model.

## Discussion

We here developed a simple empirical model predicting neural tissue stiffness based on a multiple linear regression analysis of AFM-based elasticity maps and resolution-matched immunostaining of cell nuclei, MBP, and GFAP intensities. The model captures the mechanical heterogeneities found in healthy spinal cord and can be easily extended to pathologically altered tissue and other tissue types.

The density of cell nuclei has been found to positively corelate with tissue stiffness in mouse spinal cord tissue (6), ruminant retinae (27), and in *Xenopus* brains (1, 28). Furthermore, enhancing neurogenesis and thus CNS cell density increased brain tissue stiffness (29), while loss of CNS cells during physiological aging or Alzheimer’s disease both correlated with reduced tissue stiffness (30, 31). Also, blocking cell proliferation in *Xenopus* embryos reduced cell densities and tissue stiffness if compared to control brains (28). Together, these studies strongly suggest that the density of cell nuclei directly regulates tissue stiffness.

Previous studies showed that myelin content correlates positively (23) and demyelination negatively (24, 32) with tissue stiffness. Our data are in line with these results, although they suggest a co-dependency on GFAP intensity. Furthermore, our results are in agreement with a recent report showing that GFAP intensity scales with tissue softening in cortical and spinal cord tissue after injuries (8). This tissue-scale softening might at least partly be attributed to a softening of reactive glial cells (33), which are characterized by an increase in GFAP expression levels. However, in Muller glial cells in the retina an increase in GFAP levels correlates with an increase in cell stiffness (34), suggesting that further studies are needed to illuminate the role of intermediate filaments and glial cell activity in regulating local tissue stiffness.

The constant *e* in Equation 1 likely contains contributions from hydrostatic pressure and the extracellular matrix (ECM), which in the CNS is comparatively homogeneously distributed (35). Thus, while these parameters may influence CNS mechanics in general, they are less likely to contribute to local heterogeneities in tissue stiffness. The overall stiffness of different tissue types strongly depends on their collagen I content (36). In the CNS, collagen I levels are very low (37), explaining why brain and spinal cord tissues are comparatively soft.

We here present a simple approach to estimate local tissue stiffness using immunohistochemistry in the adult mouse spinal cord. A combination of immunofluorescence images of cell densities, myelination, and glial cell content and activity was sufficient to predict local mechanical heterogeneities. Future studies will show which of these parameters are causally linked to tissue stiffness and how, and if further parameters contribute to setting up mechanical heterogeneities. Similar approaches as presented here can be taken to estimate the mechanical properties of CNS tissue during pathological processes, and of other tissue types, thus minimizing the need of complex tissue mechanics measurements and facilitating mechanobiological studies.

## Acknowledgments

The authors would like to thank Jolanta Kozlowski, Kathrin Holtzmann, Giovanna Pommerschein, Sofie Groth and Alex Winkel (JPK) for discussions and technical help. This work was supported by the Köln Fortune Program/Faculty of Medicine, University of Cologne (Fellowship to D.E.K.), German National Academic Foundation (Scholarship to D.E.K.), Herchel Smith Foundation (Fellowship to E.M.), Deutsche Forschungsgemeinschaft (grant KU2760/2-1 to S.K.), UK Medical Research Council (Career Development Award G1100312/1 to K.F.), and the Human Frontier Science Program (Young Investigator Grant RGY0074/2013 to K.F.).

## Author contributions

D.E.K. and K.F. conceived and designed the study. D.E.K. performed all experiments and analysed the data. E.M. helped with automatization of the analysis and S.K. helped with immunohistochemistry. S.K. and K.F. supervised the work. D.E.K. and K.F. wrote the paper with contributions from all authors.

